## Supplementary Figures for "High fat diet ameliorates mitochondrial cardiomyopathy in CHCHD10 mutant mice"

### Supplementary Fig. 1

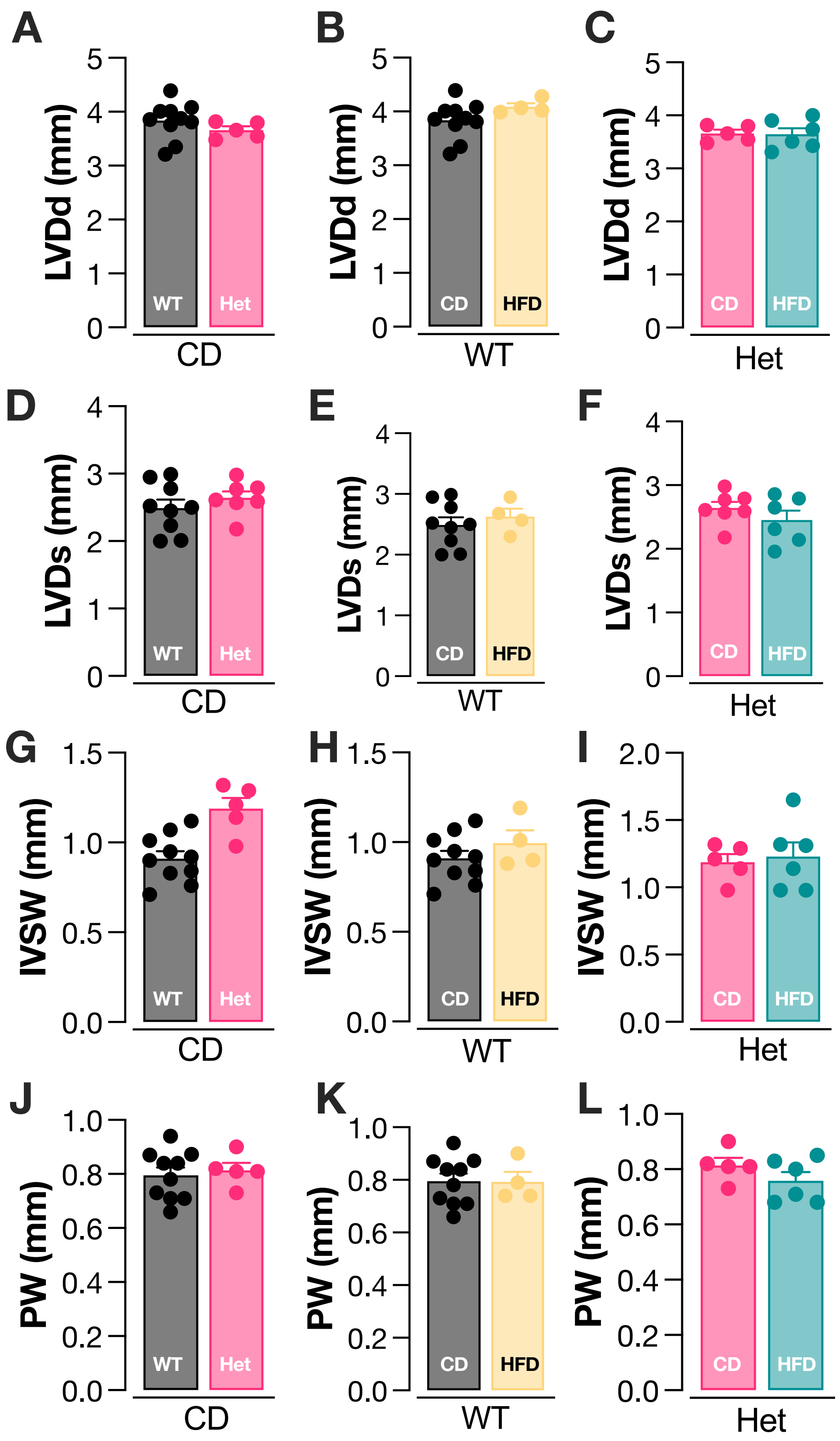

#### Supplementary Fig. 2

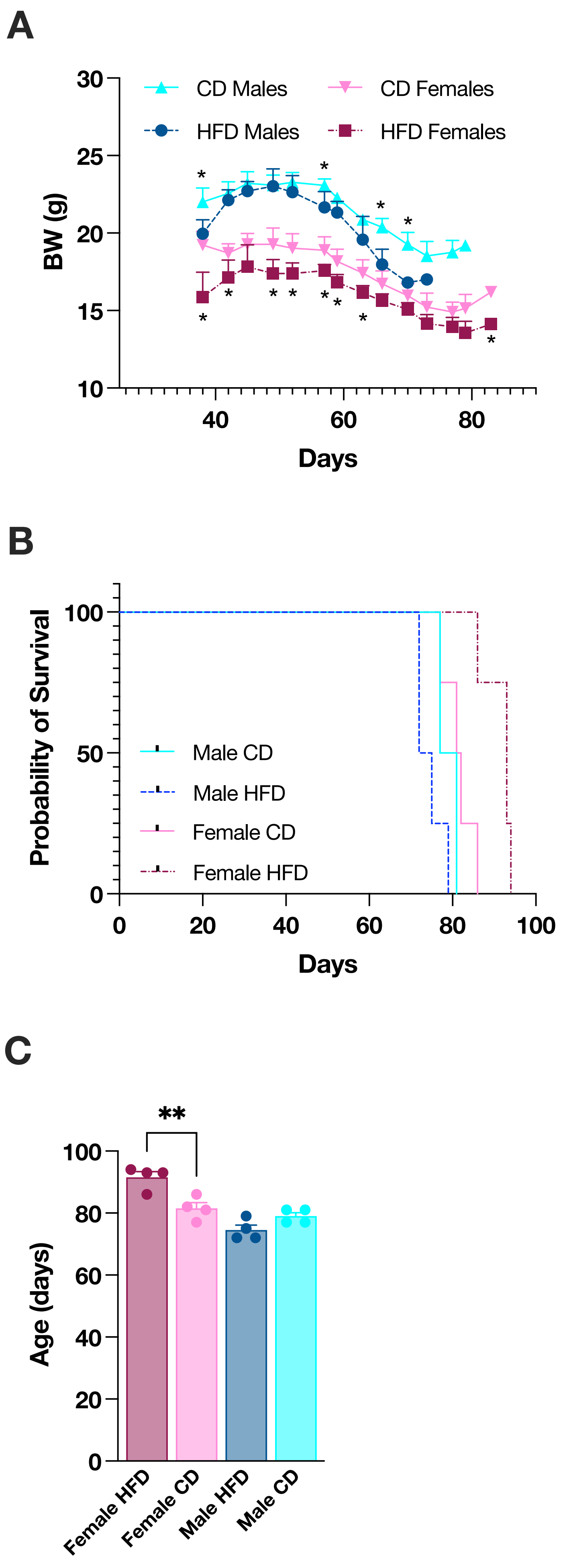

### Supplementary Fig. 3

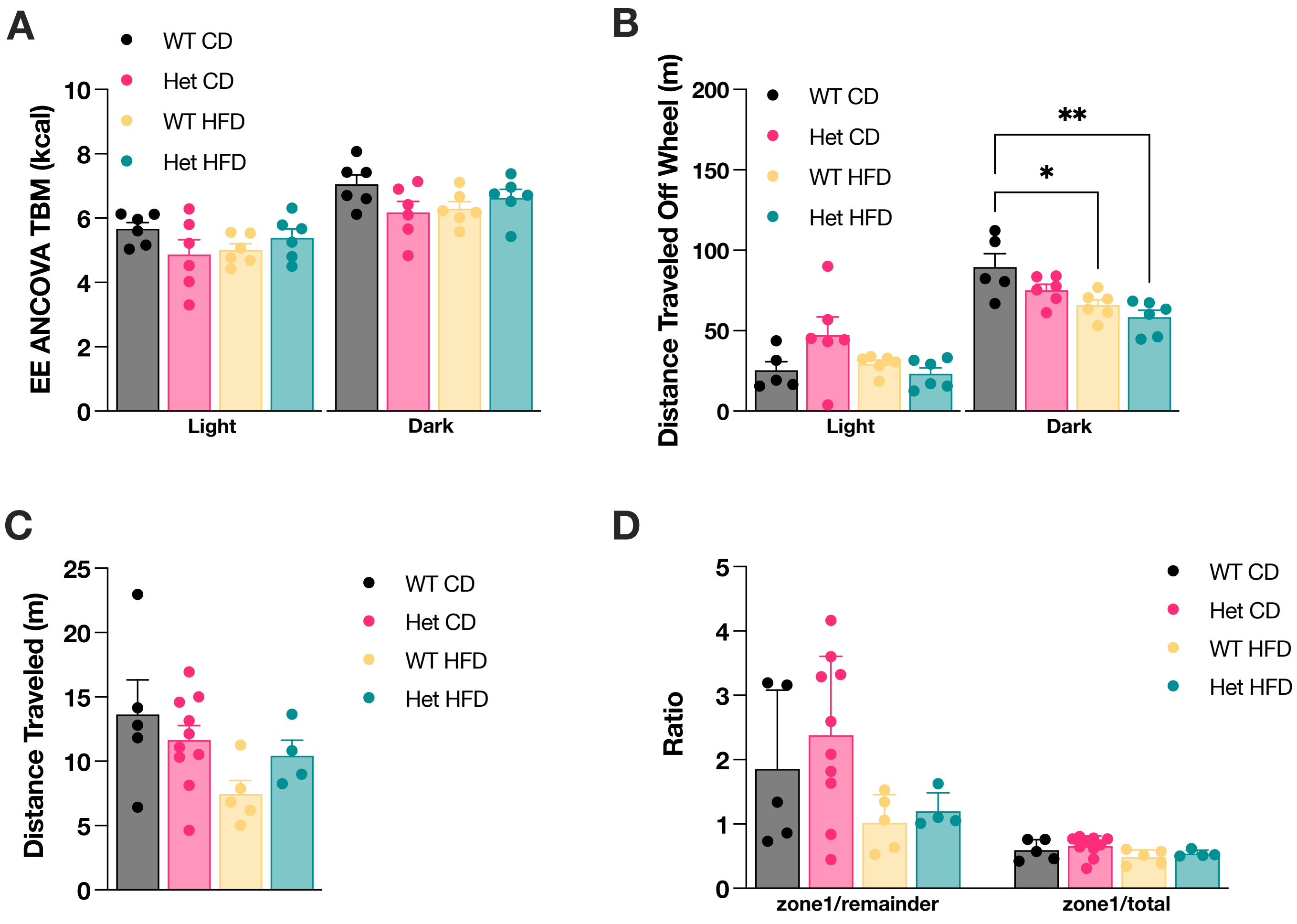

### Supplementary Fig. 4

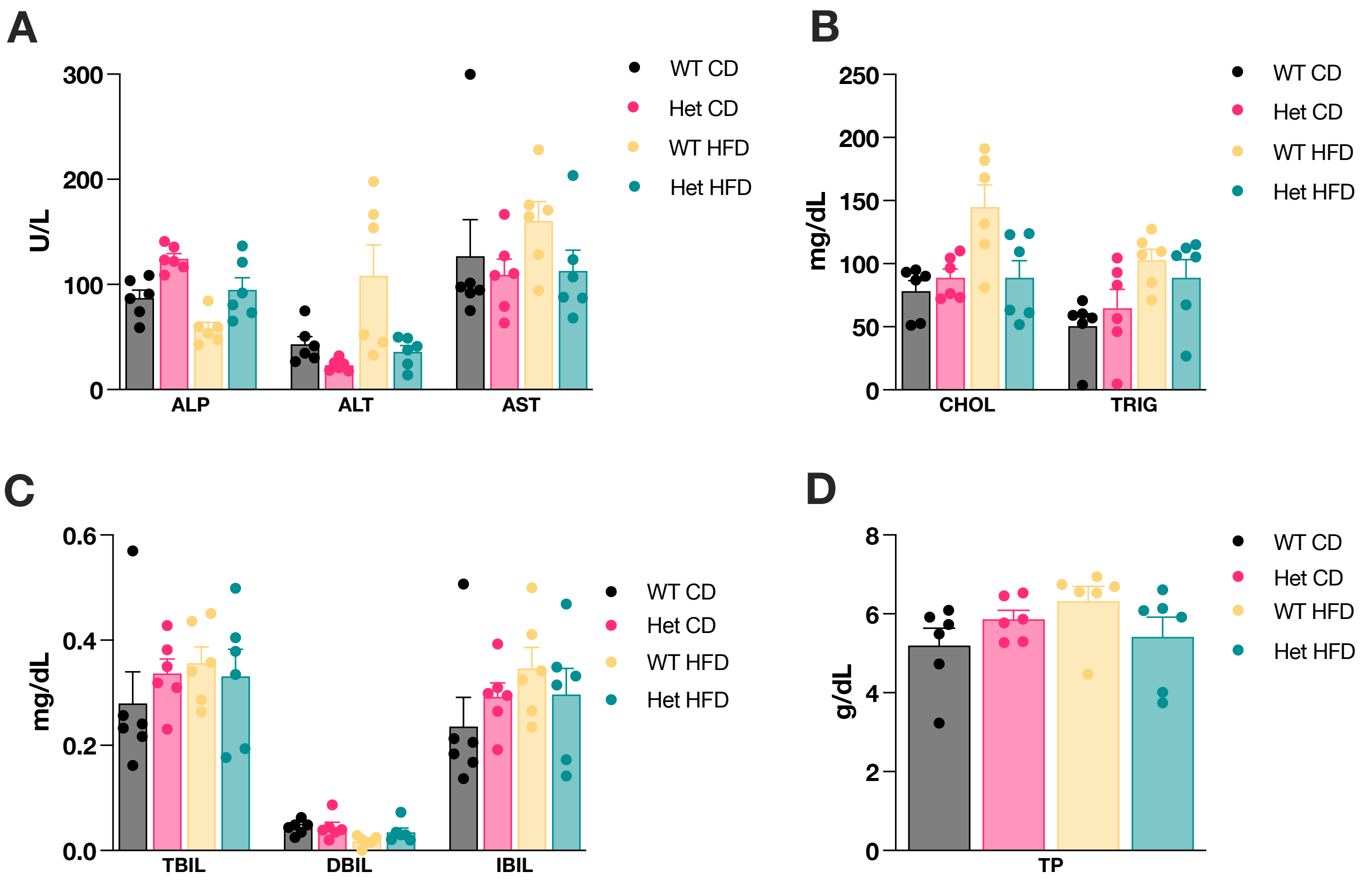

### Supplementary Fig. 5

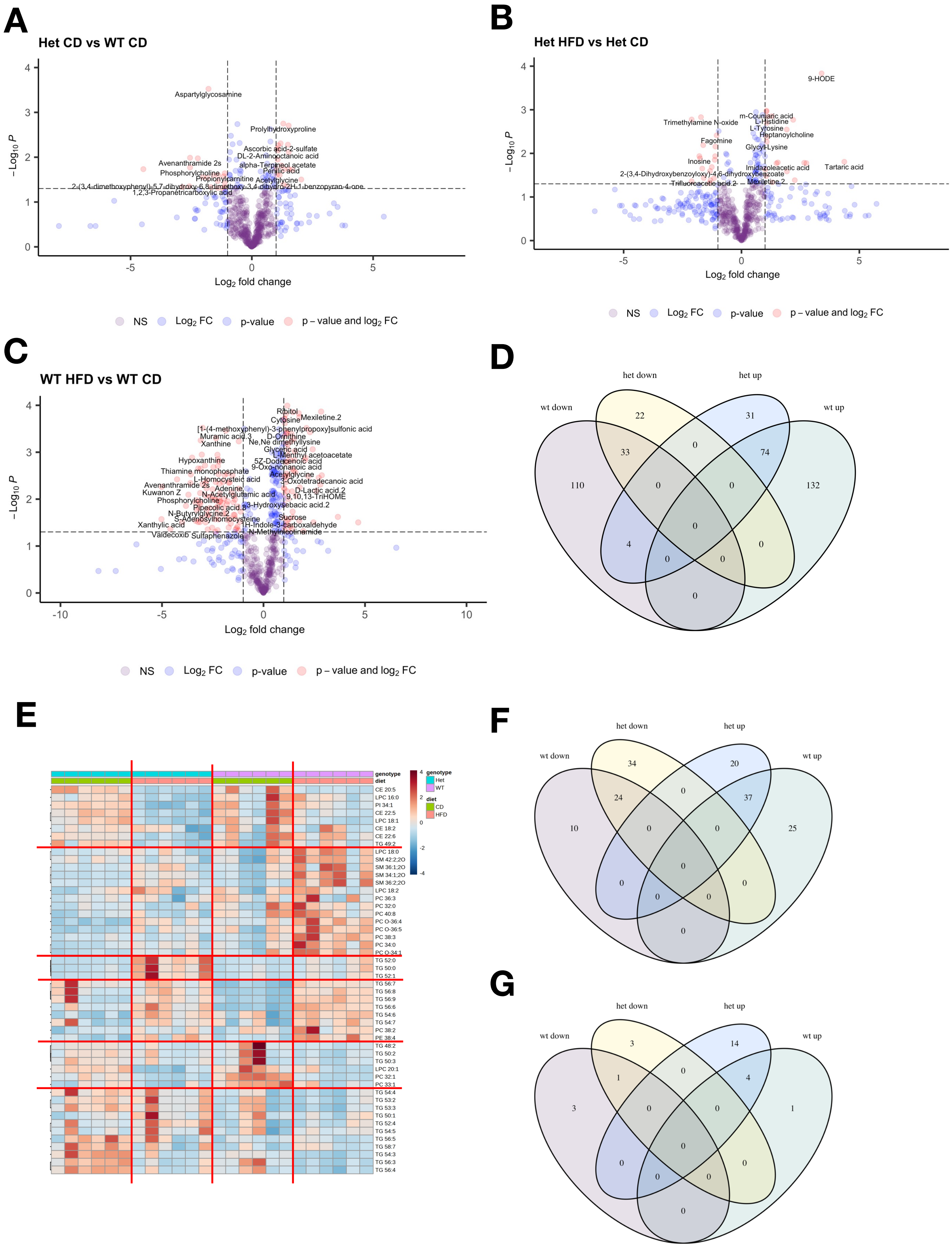

### Supplementary Fig. 6

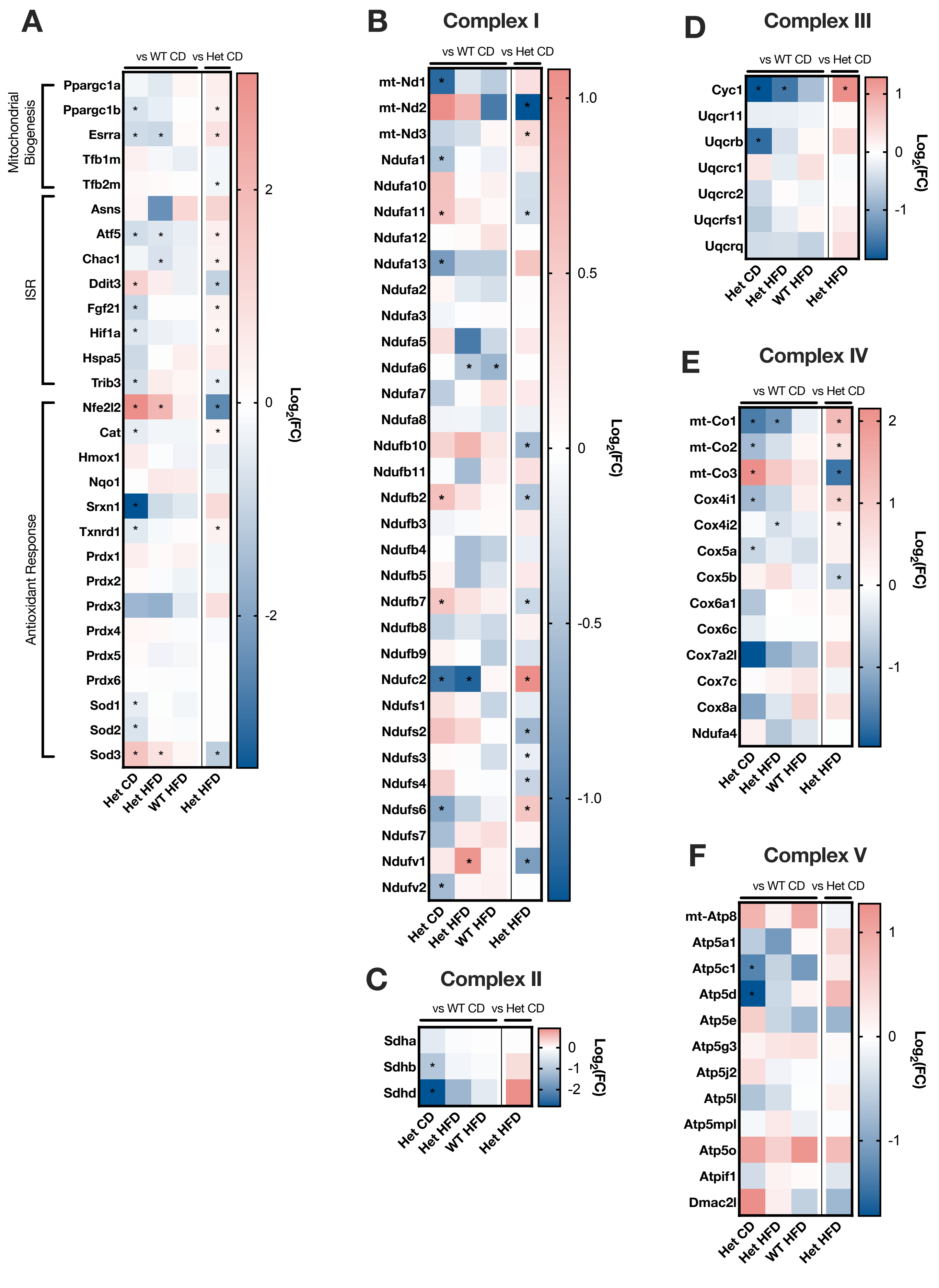

### Supplementary Fig. 7

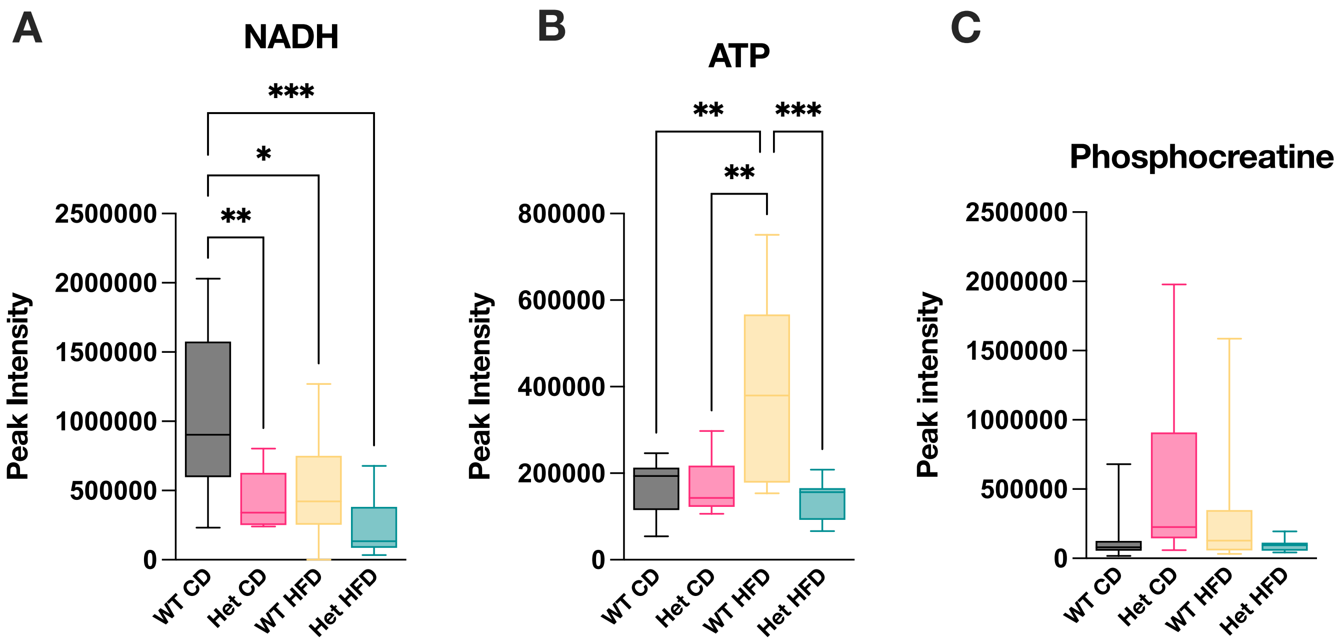

### Supplementary Fig. 8

A

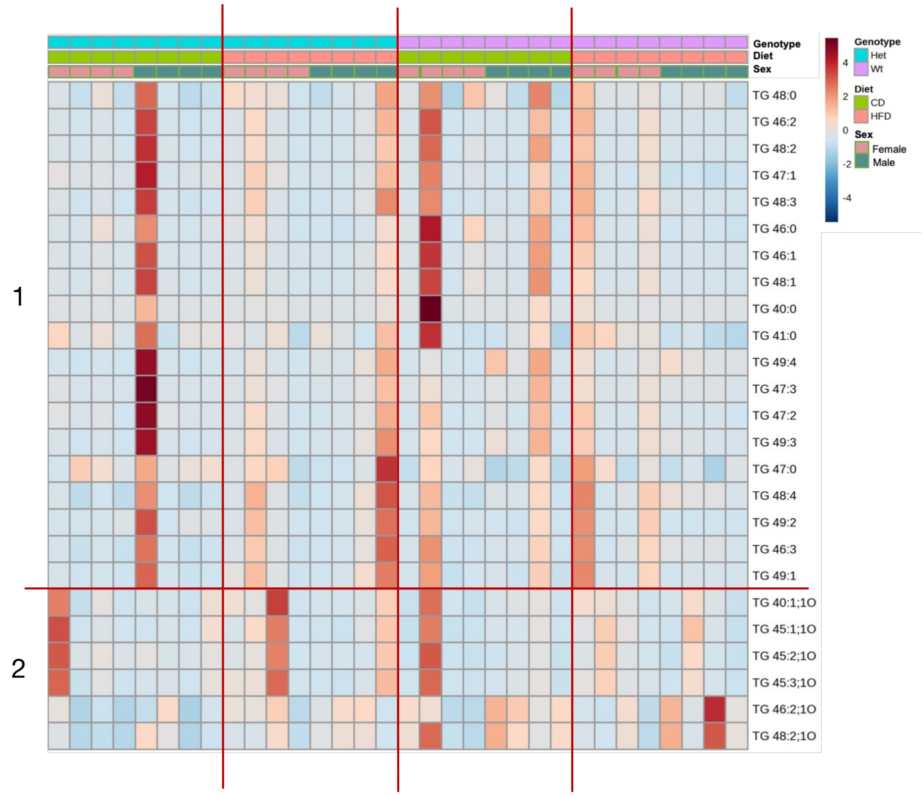


B

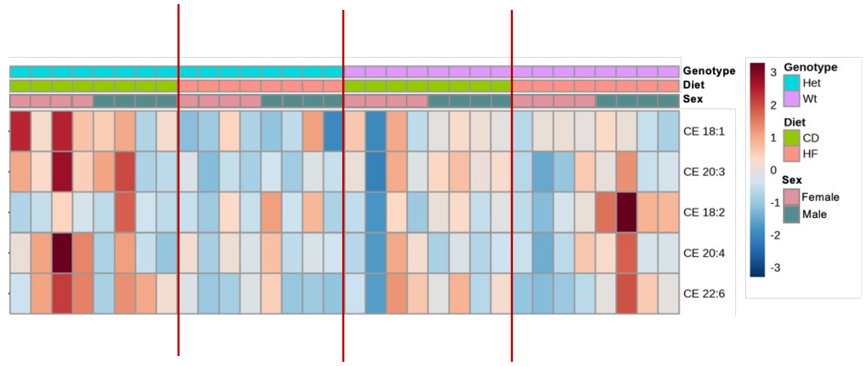
